# GCalignR: An R package for aligning Gas-Chromatography data

**DOI:** 10.1101/110494

**Authors:** Meinolf Ottensmann, Martin A. Stoffel, Hazel J. Nichols, Joseph I. Hoffman

**Author notes:** Joint first authors (Meinolf Ottensmann, Martin A. Stoffel).

## Abstract

Chemical cues are arguably the most fundamental means of animal communication and play an important role in mate choice and kin recognition. Consequently, gas chromatography (GC) in combination with either mass spectrometry (MS) or flame ionisation detection (FID) are commonly used to characterise complex chemical samples. Both GC-FID and GC-MS generate chromatograms comprising peaks that are separated according to their retention times and which represent different substances. Across chromatograms of different samples, homologous substances are expected to elute at similar retention times. However, random and often unavoidable experimental variation introduces noise, making the alignment of homologous peaks challenging, particularly with GC-FID data where mass spectral data are lacking. Here we present `GCalignR`, a user-friendly R package for aligning GC-FID data based on retention times. The package also implements dynamic visualisations to facilitate inspection and fine-tuning of the resulting alignments, and can be integrated within a broader workflow in R to facilitate downstream multivariate analyses. We demonstrate an example workflow using empirical data from Antarctic fur seals and explore the impact of user-defined parameter values by calculating alignment error rates for multiple datasets. The resulting alignments had low error rates for most of the explored parameter space and we could also show that `GCalignR` performed equally well or better than other available software. We hope that `GCalignR` will help to simplify the processing of chemical datasets and improve the standardization and reproducibility of chemical analyses in studies of animal chemical communication and related fields.

## Introduction

Chemical cues are arguably the most common mode of communication among animals [1]. In the fields of animal ecology and evolution, increasing numbers of studies have therefore been using approaches like gas chromatography (GC) to characterise the chemical composition of body odours and scent marks. These studies have shown that a variety of cues are chemically encoded, including phylogenetic relatedness [2], breeding status [3], kinship [4–6] and genetic quality [6–8].

GC vaporises a chemical sample and retards its components differentially based on their chemical properties while passing a gas through a column. The chemical composition of the sample can then be resolved using a number of approaches such as GC coupled to a flame ionization detector (GC-FID) or GC coupled to a mass spectrometer (GC-MS). GC-FID produces a chromatogram in which each substance is represented by a peak, the area of which is proportional to the concentration of that substance in the sample [9]. Although GC-FID is a relatively inexpensive and high-throughput approach, the substances themselves can only be characterised according to their retention times, so their chemical composition remains effectively unknown. GC-MS similarly generates a chromatogram, but additionally provides spectral profiles corresponding to each peak, thereby allowing putative identification by comparison to databases of known substances. Both approaches have distinct advantages and disadvantages, but the low cost of GC-FID, coupled with the fact that most chemicals in non-model organisms do not reveal matches to databases containing known chemicals, has led to an increasing uptake of GC-FID in studies of wild populations [10–13]. GC-FID is particularly appropriate for studies seeking to characterise broad patterns of chemical similarity without reference to the exact nature of the chemicals involved.

As a prerequisite for any downstream analysis, homologous substances across samples need to be matched. Therefore, an important step in the processing of the chemical data is to construct a so called peak list, a matrix containing the relative abundances of each homologous substance across all of the samples. With GC-MS, homologous substances can be identified on the basis of both their retention times and the accompanying spectral information. However, with GC-FID, homologous substances can only be identified based on their retention times. This can be challenging because these retention times are often perturbed by subtle, random and often unavoidable experimental variation including changes in ambient temperature, flow rate of the carrier gas and column ageing [14, 15].

Numerous algorithms have been developed for aligning MS data (reviewed by [16]). Although these are all capable of aligning homologous peaks within chromatograms, most of them rely on the use of spectral information provided by MS. Consequently, the majority of software that implement these algorithms do not support GC-FID data due to the lack of spectral information. A handful of these programs are able to work with a peak list format (e.g. amsrpm [17] and `ptw` [18]) but the underlying algorithms may not be well suited to GC-FID data for two main reasons. First, the alignment is conducted strictly pairwise with respect to a pre-defined reference sample, because in general the focus is on a relatively small pool of substances that are expected to be present in most if not all samples [19]. However, applied to wild animal populations, GC-FID often yields high diversity datasets in which only a small subset of chemicals may be common to all individuals [6, 20]. Second, existing algorithms are known to be sensitive to variation in peak intensity, which is expected in GC-FID datasets and may contain important biological information [6, 20–22].

To tackle the above issues, a program called GCALIGNER was recently written in Java for aligning GC-FID data [23]. This program appears to perform well based on three test datasets, each corresponding to a different bumblebee species (`Bombus` spp.). However, the underlying algorithm compares each peak with the following peak in the same sample and therefore cannot align the last peak [23]. Moreover, with the increasing popularity of open source environments such as R, there is a growing need for software that can be easily integrated into broader workflows, where the source code can be modified and potentially further extended by the user, and where related tools like Rmarkdown [24] can be applied to maximise transparency and reproducibility [25]. Furthermore, especially for GC-FID data where spectral data are not available, a useful addition would be to integrate dynamic visualisation tools into software to facilitate the evaluation and subsequent fine-tuning of alignment parameters.

In order to determine which alignment tools are commonly used in the fields of ecology and evolution, we conducted a bibliographic survey, focusing on the journals ‘Animal Behaviour’ and ‘Proceedings of the Royal Society B’, which recovered a total of 38 studies using GC-FID or GC-MS to investigate scent profiles (see S1 for details). None of these studies used any form of alignment tool but rather aligned and called the peaks manually (e.g. [26]), a time-consuming process that can be prone to bias [27] and detrimental to reproducibility.

To address the above issues, we developed `GCalignR`, an R package for aligning GC-FID data, but which can also align any type of GC data where peaks are characterised by retention times. The package implements a fast and objective method to cluster putatively homologous substances prior to multivariate statistical analyses. Using sophisticated visualisations, the resulting alignments can then be fine tuned. Finally, the package provides a seamless transition from the processing of the peak data through to downstream analysis within other widely used R packages for multivariate analysis, such as vegan [28].

In this paper, we present `GCalignR` and describe the underlying algorithms and their implementation within a suite of R functions. We provide an example workflow using a previously published chemical dataset of Antarctic fur seals (*Arctocephalus gazella*) that shows a clear distinction between animals from two separate breeding colonies [6]. We then compare the performance of `GCalignR` with GCALIGNER based on the same three bumblebee datasets given in [23] and explore the sensitivity of `GCalignR` to user-defined alignment parameter values. Finally, we compared our alignment procedure with a very different approach –parametric time warping– which is commonly used in the fields of proteomics and metabolomics [18, 38].

## Material and methods

### Overview of the package

Fig 1 shows an overview of `GCalignR` in the context of a workflow for analysing GC-FID data within R. A number of steps are successively implemented, from checking the raw data through aligning peak lists and inspecting the resulting alignments to normalising the peak intensity measures prior to export into vegan [28]. In brief, the alignment procedure is implemented in three consecutive steps that start by accounting for systematic shifts in retention times among samples and subsequently align individual peaks based on variation in retention times across the whole dataset. For simplicity, this procedure is embedded within a single function (`align_chromatograms`) that allows the customisation of peak alignments by adjusting a combination of three parameters. The package vignettes provide a detailed description of all of the functions and their arguments and can be accessed via `browseVignettes(‘GCalignR’)` after the package has been installed.

**Fig 1.**
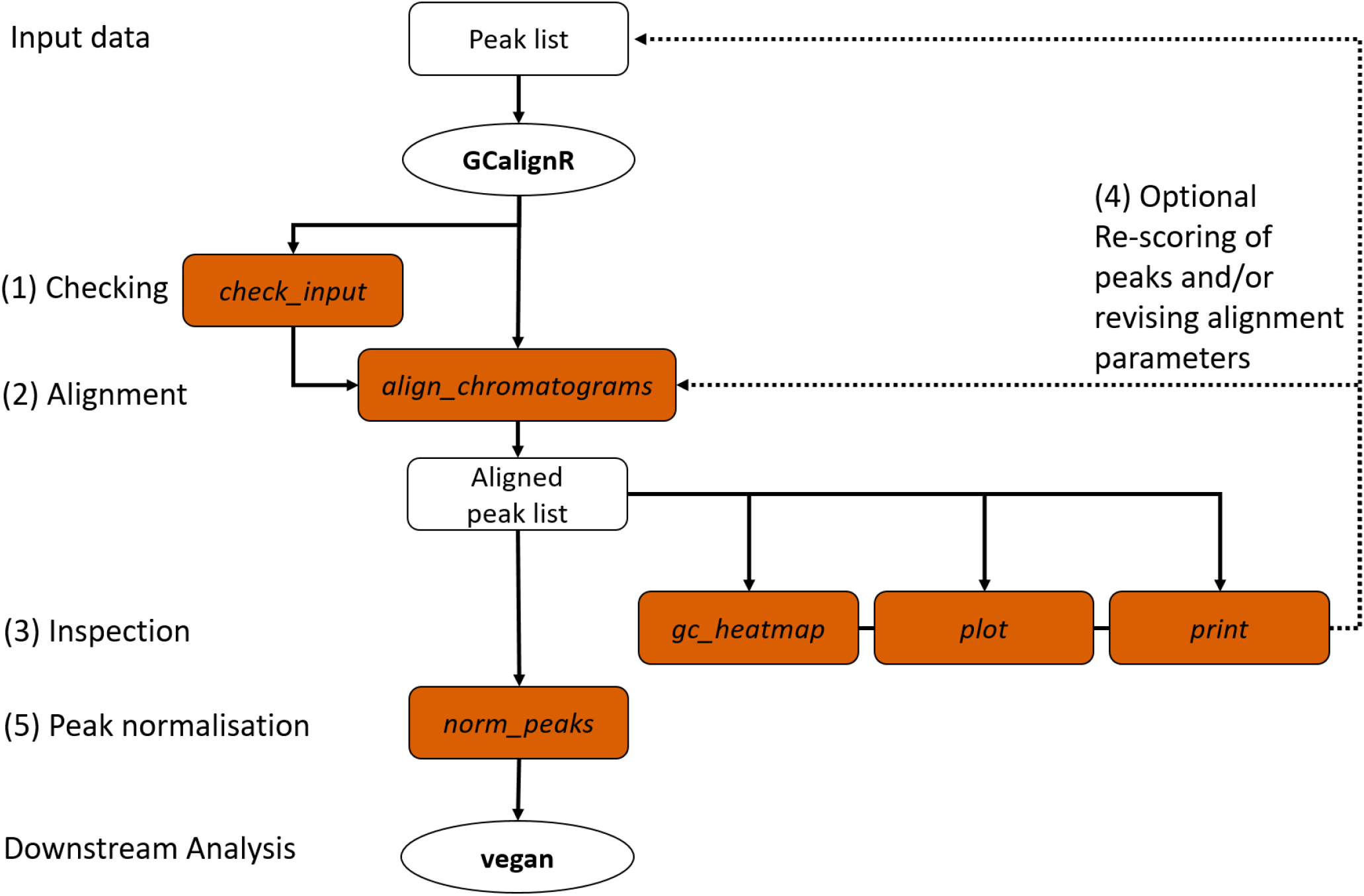
Overview of the GCalignR workflow. The steps listed in the main text are numbered from one to five and the filled boxes represent functions of the package (see Materials and methods for details).

### Raw data format and conversion to working format

GC-FID produces raw data in the form of individual chromatograms that show the measured electric current over the time course of a separation run. Proprietary software provided by the manufacturers of GC-FID machines (e.g. ‘LabSolutions’, Shimadzu; ‘Xcalibur’, Thermo Fisher and ‘ChemStation’, Agilent Technologies) are then used to integrate and export peaks in the format of a table containing retention times and intensity values (e.g. peak area and height). Fig 2 A shows chromatograms of three hypothetical samples where peaks have been integrated and annotated with retention times and peak heights. The corresponding input format comprising a table of retention times and peak heights is also shown. The working format of `GCalignR` is a retention time matrix in which each sample corresponds to a column and each peak corresponds to a row (see Fig 2 B).

**Fig 2.**
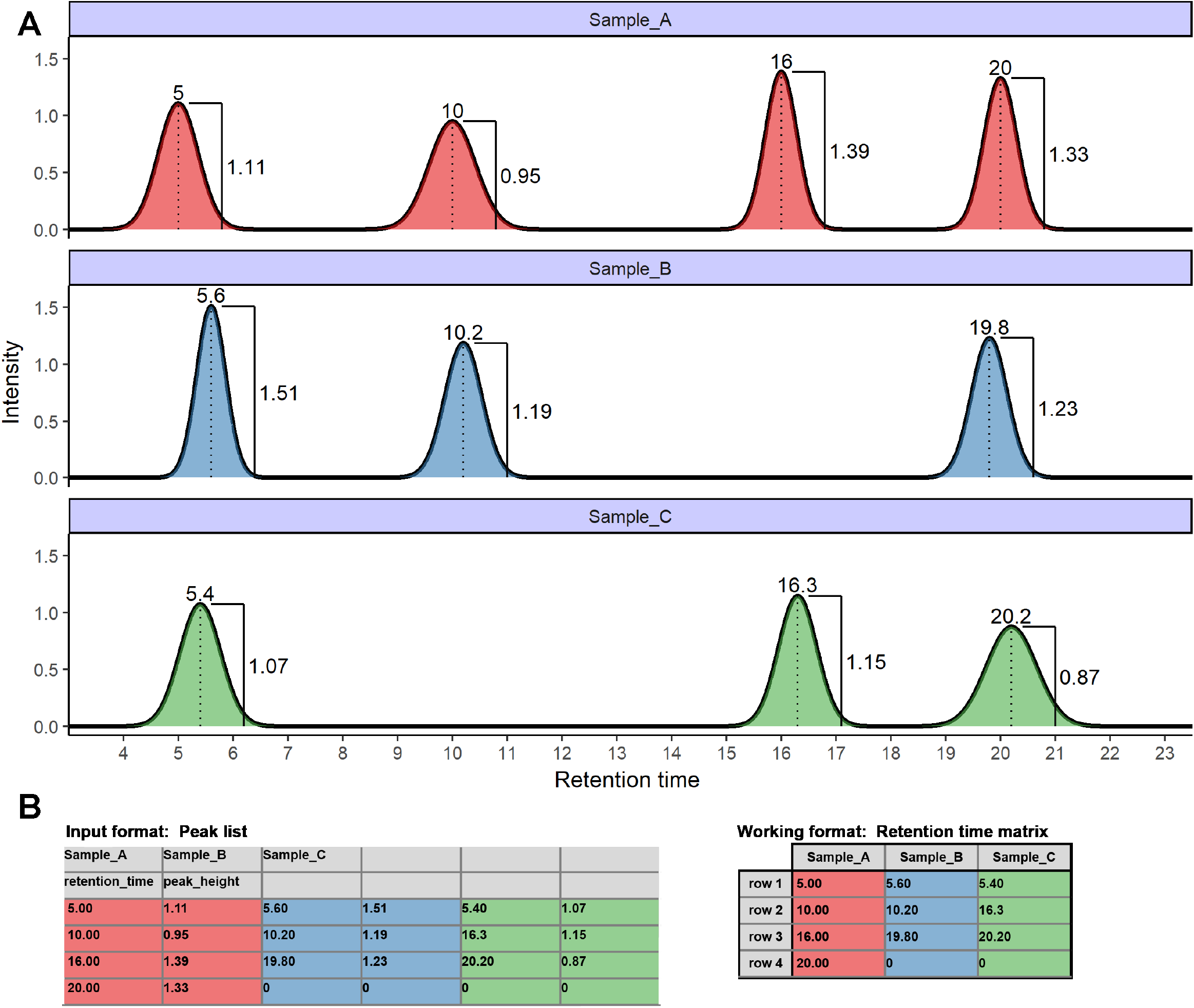
GC-FID data formats. **A** Three hypothetical chromatograms are shown corresponding to samples A, B and C. Integrated peaks (filled areas) are annotated with retention times and peak heights. **B** Using proprietary software (see main text), retention times and quantification measures like the peak height can be extracted and written to a peak list that contains sample identifiers (’Sample_A’, ’Sample_B’ and ’Sample_C’), variable names (’retention_time’ and ’peak_height’) and respective values. Computations described in this manuscript use a retention matrix as the working format

### Overview of the Alignment algorithm

We developed an alignment procedure based on dynamic programming [29] that involves three sequential steps to align and finally match peaks belonging to putatively homologous substances across samples (see Fig 3 for a flowchart and Fig 4 for a more detailed schematic representation). All of the raw code for implementing these steps is available via GitHub and CRAN and each step is described in detail below. The first step is to align each sample to a reference sample while maximising overall similarity through linear shifts of retention times. This procedure is often described in the literature as ’full alignment’ [18]. In the second step, individual peaks are sorted into rows based on close similarity of their retention times, a procedure that is often referred to as ’partial alignment’ [18]. Finally, there is still a chance that homologous peaks can be sorted into different, but adjacent, rows in different samples, depending on the variability of their retention times (for empirical examples, see S2, Fig 5 and 11). Consequently, a third step merges rows representing putatively homologous substances.

**Fig 3.**
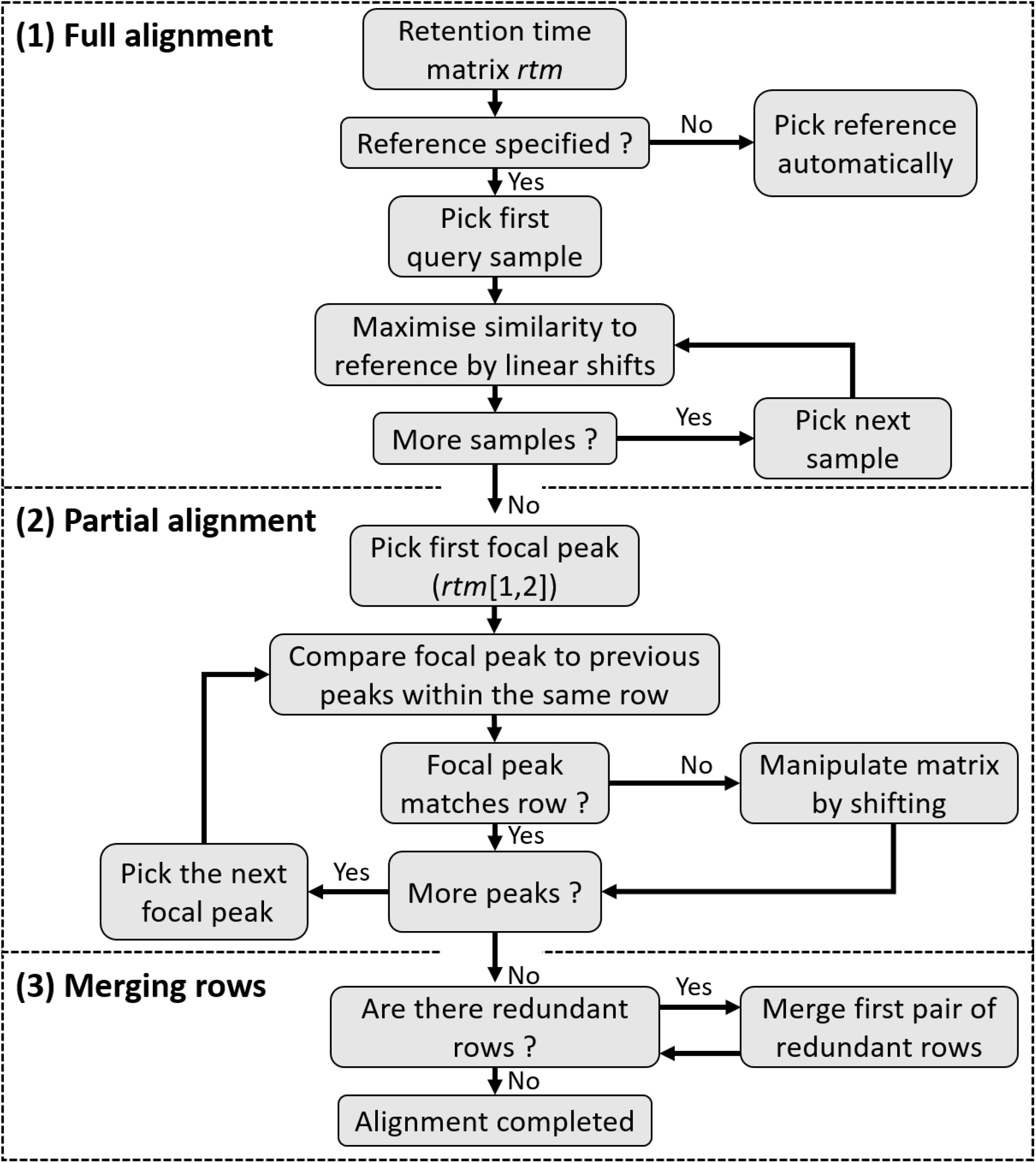
A flow chart showing the three sequential steps of the alignment algorithm of the peak alignment method.

**Fig 4.**
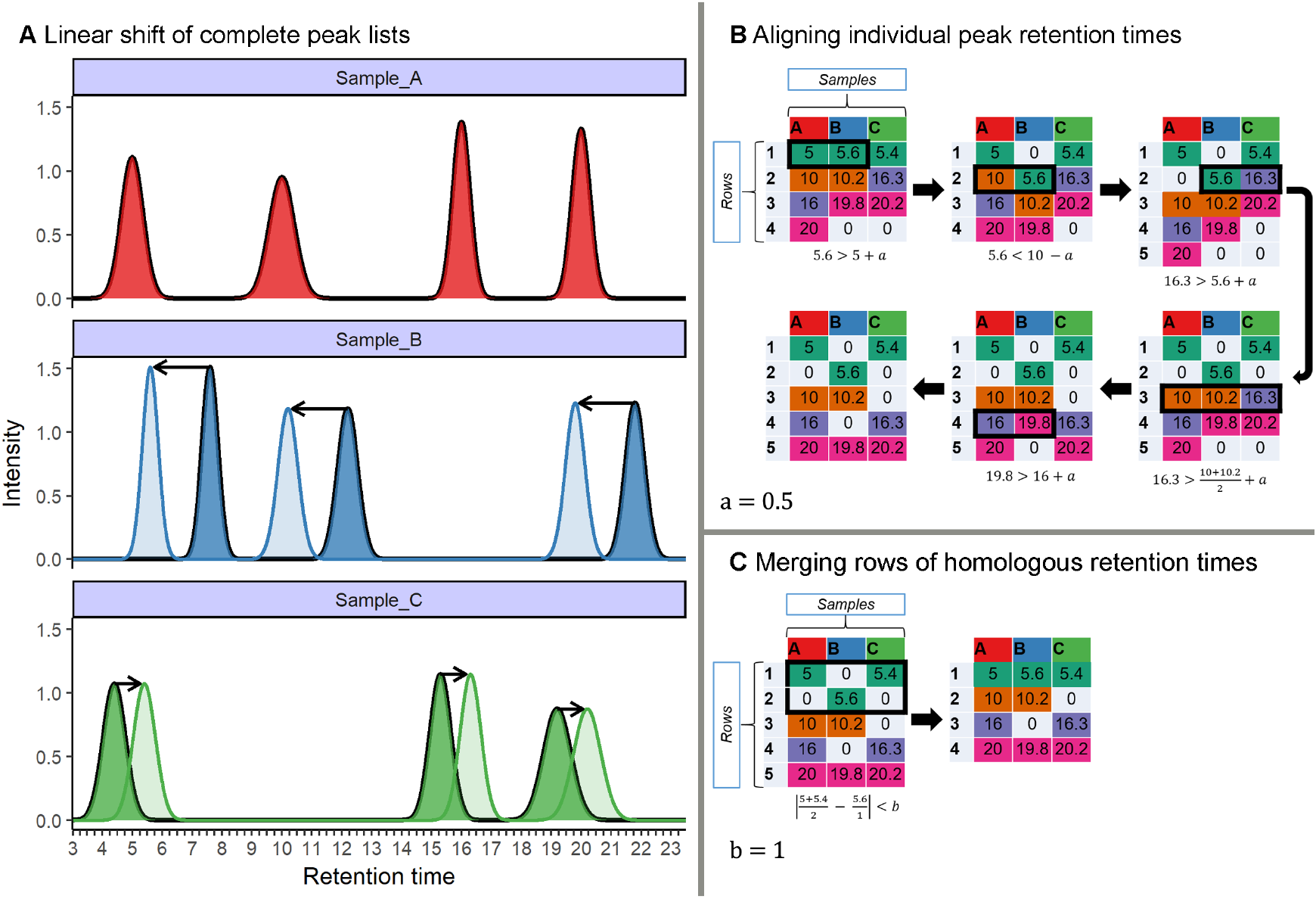
Overview of the three-step alignment algorithm implemented in GCalignR using a hypothetical dataset. **A.** Linear shifts are implemented to account for systematic drifts in retention times between each sample and the reference (Sample_A). In this hypothetical example, all of the peaks within Sample_B are shifted towards smaller retention times, while the peaks within Sample_C are shifted towards larger retention times. **B** and **C** work on retention time matrices, in which rows correspond to putative substances and columns correspond to samples. For illustrative purposes, each cell is colour coded to refer to the putative identity of each substance in the final alignment. **B.** Consecutive manipulations of the matrices are shown in clockwise order. Here, black rectangles indicate conflicts that are solved by manipulations of the matrices. Zeros indicate absence of peaks and are therefore not considered in computations. Peaks are aligned row by row according to a user-defined criterion, *a* (see main text for details). **C.** Rows of similar mean retention time are subsequently merged according to the user-defined criterion, *b* (see main text for details).

**Fig 5.**
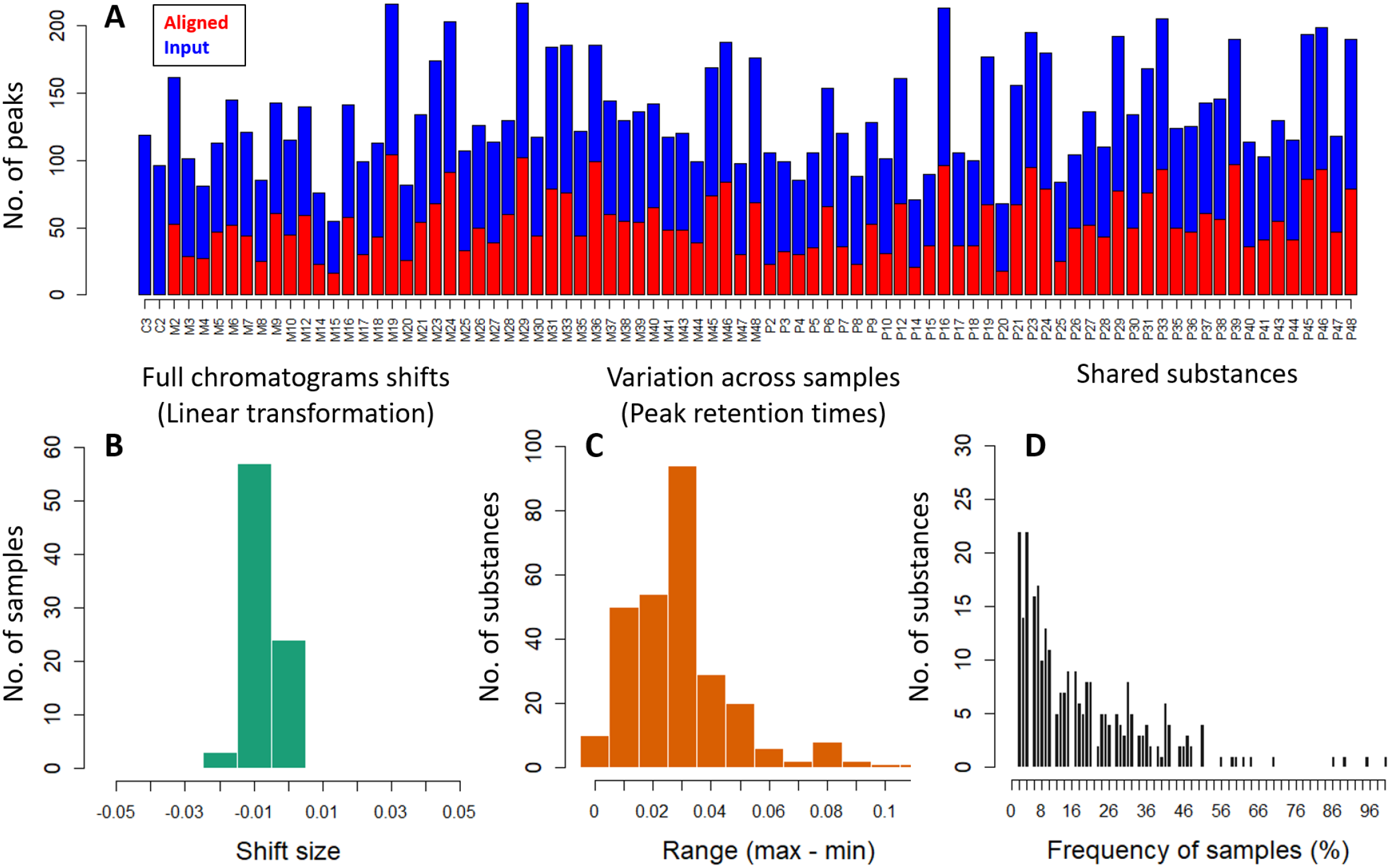
Diagnostic plots summarising the alignment of the Antarctic fur seal chemical dataset. **A** shows the number of peaks both prior to and after alignment; **B** shows a histogram of linear shifts across all samples; **C** shows the variation across samples in peak retention times; and **D** shows a frequency distribution of substances shared across samples.

#### (i) Full alignment of peaks lists

The first step in the alignment procedure consists of an algorithm that corrects systematic linear shifts between peaks of a query sample and a fixed reference to account for systematic shifts in retention times among samples (Fig 4 A). Following the approach of Daszykowski et al. [30], the sample that is most similar on average to the other samples can be automatically selected as a reference by choosing the sample with the lowest median deviation score weighted by the number of peaks to avoid a bias towards samples with few peaks:

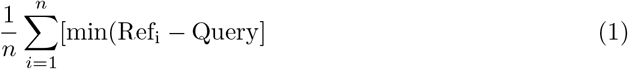

where n is the number of retention times in the reference sample. Alternatively, the reference can be specified by the user. Using a simple warping method [31], the complete peak list of the query is then linearly shifted within an user-defined retention time window with an interval of 0.01 minutes. For all of the shifts, the summed deviation in retention times between each reference peak and the nearest peak in the query is used to approximate similarity as follows:

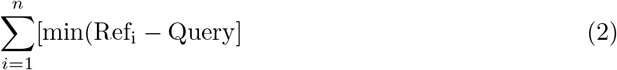

where n is the number of retention times in the reference sample. With increasing similarity, this score will converge towards zero the more homologous peaks are aligned, whereas peaks that are unique to either the query or the reference are expected to behave independently and will therefore have little effect on the overall score. The shift yielding to the smallest score is selected to transform retention times for the subsequent steps in the alignment (Fig 4 B and C). As the effectiveness of this approach relies on a sufficient number of homologous peaks that can be used to detect linear drift, the performance of the algorithm may vary between datasets.

#### (ii) Partial alignment of peaks

The second step in the alignment procedure aligns individual peaks across samples by comparing the peak retention times of each sample consecutively with the mean of all previous samples (Fig 4 B) within the same row. If the focal cell within the matrix contains a retention time that is larger than the mean retention time of all previous cells within the same row plus a user-defined threshold (Eq (3)), that cell is moved to the next row.

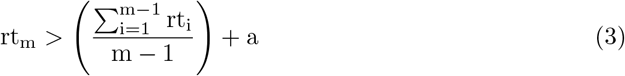

where rt is the retention time; m is the focal cell and a is the user-defined threshold deviation from the mean retention time. If the focal cell contains a retention time that is smaller than the mean retention time of all previous cells within the same row minus a user-defined threshold (Eq (4)), all previous retention times are then moved to the next row.

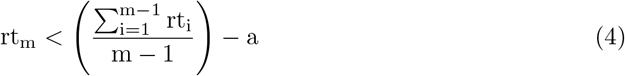

After the last retention time of a row has been evaluated, this procedure is repeated for the next row until the end of the retention time matrix is reached (Fig 4 B).

#### (iii) Merging rows

The third step in the alignment procedure accounts for the fact that a number of homologous peaks will be sorted into multiple rows that can be subsequently merged (Fig 4 C). However, this results in a clear pattern whereby some of the samples will have a retention time in one of the rows while the other samples will have a retention time in an adjacent row (S2, Fig 5 and 11). Consequently, pairs of rows can be merged when this does not cause any loss of information, an assumption that is true as long as no sample exists that contains peaks in both rows, (Fig 4 C). The user can define a threshold value in minutes (i.e. parameter b in Fig 4 C) that determines whether or not two such adjacent rows are merged. While the described pattern is unlikely to occur in large datasets purely by chance for non-homologous peaks, small datasets may require more strict threshold values to be selected.

### Implementation of the alignment method

The alignment algorithms that are described above are all executed by the core function `align_chromatograms` based on the user-defined parameters shown in (Table 1). Of these, parameters (`max_linear_shift, max_diff_peak2mean` and `min_diff_peak2peak`) can be adjusted by the user to fine-tune the alignment procedure. There a several additional parameters that allow for optional processing and filtering of the data independently of the alignment procedure. For further details, the reader is referred to the accompanying vignettes (see S3 and S4) and helpfiles of the R package.

**Table 1.**
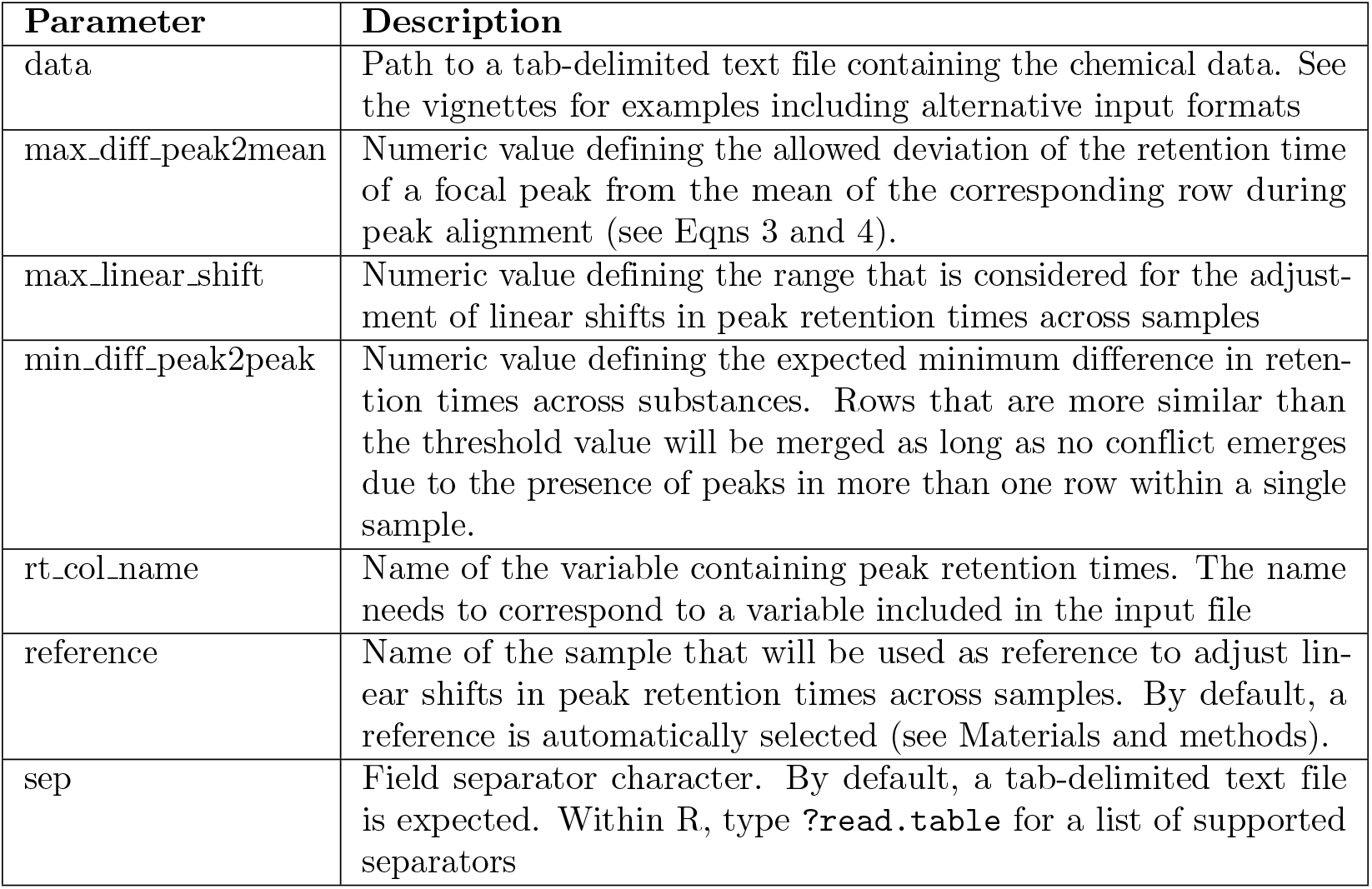
Mandatory arguments of the function align_chromatograms.

### Demonstration of the workflow

Here, we demonstrate a typical workflow in `GCalignR` using chemical data from skin swabs of 41 Antarctic fur seal (*Arctocephalus gazella*) mother-pup offspring pairs from two neighbouring breeding colonies at South Georgia in the South Atlantic. Sample collection and processing are described in detail in Stoffel et al. [6]. In brief, chemical samples were obtained by rubbing the cheek, underneath the eye, and behind the snout with a sterile cotton wool swab and preserved in ethanol stored prior to analysis. In order to account for possible contamination, two blank samples (cotton wool with ethanol) were processed and analysed using the same methodology. Peaks were integrated using Xcalibur (Thermo Scientific). The chemical data associated with these samples are provided in the file `peak_data.txt`, which is distributed together with `GCalignR`. Additional data on colony membership and age-class are provided in the data frame `peak_factors.RData`.

Prior to peak alignment, the `check_input` function interrogates the input file for typical formatting errors and missing data. We encourage the use of unique names for samples consisting only of letters, numbers and underscores. If the data fail to pass this quality test, indicative warnings will be returned to assist the user in error correction. As this function is executed internally prior to alignment, the data need to pass this check before the alignment can begin.

~~~
# load the package
library (GCalignR)
# set the path to the input data
fpath <- system.file (dir = “extdata”,
                            file = “peak_data.txt”,
                            package = “GCalignR”)
# check for formatting problems
check_input (fpath)
~~~

In order to begin the alignment procedure, the following code needs to be executed:

~~~
aligned_peak_data <- align_chromatograms(data = peak_data,
         rt_col_name = “ time “,
         max_diff_peak2mean = 0.02,
         min_diff_peak2peak = 0.08,
         max_linear_shift = 0.05,
         delete_single_peak = TRUE,
         blanks = c(“C2”, ”C3”))
~~~

Here, we set `max_linear_shift` to 0.05, `max_diff_peak2mean` to 0.02 and `min_diff_peak2peak` to 0.08. By defining the argument `blanks`, we implemented the removal of all substances that are shared with the negative control samples from the aligned dataset. Furthermore, substances that are only present in a single sample were deleted from the dataset using the argument `delete_single_peak = TRUE` as these are not informative in analysing similarity pattern [32]. Afterwards, a summary of the alignment process can be retrieved using the printing method, which summarises the function call including defaults that were not altered by the user. This provides all of the relevant information to retrace every step of the alignment procedure.

~~~
# verbal summary of the alignment
print (aligned_peak_data)
~~~

As alignment quality may vary with the parameter values selected by the user, the plot function can be used to output four diagnostic plots. These allow the user to explore how the parameter values affect the resulting alignment and can help to flag issues with the raw data.

~~~
# produces Fig. 5
plot (aligned_peak_data)
~~~

The resulting output for the Antarctic fur seal chemical dataset, shown in Fig 5, reveals a number of pertinent patterns. Notably, the removal of substances shared with the negative controls or present in only one sample resulted in a substantial reduction in the total number of peaks present in each sample (Fig 5 A). Furthermore, for the majority of the samples, either no linear shifts were required, or the implemented transformations were very small compared to the allowable range (Fig 5 B). Additionally, the retention times of putatively homologous peaks in the aligned dataset were left-skewed, indicating that the majority of substances vary by less than 0.05 minutes (Fig 5 C) but there was appreciable variation in the number of individuals in which a given substance was found (Fig 5 D).

Additionally, the aligned data can be visualised using a heat map with the function `gc_heatmap`. Heat maps allow the user to inspect the distribution of aligned substances across samples and assist in fine-tuning of alignment parameters as described within the vignettes (see S3 and S4) and S2.

~~~
gc_heatmap (aligned_peak_data)
~~~

### Peak normalisation and downstream analyses

In order to account for differences in sample concentration, peak normalisation is commonly implemented as a pre-processing step in the analysis of olfactory profiles [33–35]. The `GCalignR` function `normalise_peaks` can therefore be used to normalise peak abundances by calculating the relative concentration of each substance in a sample. The abundance measure (e.g. peak area) needs to be specified as `conc_col_name` in the function call. By default, the output is returned in the format of a data frame that is ready to be used in downstream analyses.

~~~
# extract normalised peak area values
scent <- norm_peaks(data = aligned_peak_data,
           rt_col_name = “time”,
           conc_col_name = “area”,
           out = “data.frame”)
~~~

The output of `GCalignR` is compatible with other functionalities in R, thereby providing a seamless transition between packages. For example, downstream multivariate analyses can be conducted within the package vegan [28]. To visualise patterns of chemical similarity within the Antarctic fur seal dataset in relation to breeding colony membership, we used non-metric-multidimensional scaling (NMDS) based on a Bray-Curtis dissimilarity matrix in vegan after normalisation and log-transformation of the chemical data.

~~~
# log + 1 transformation
scent <- log(scent + 1)
# sorting by row names
scent <- scent [match(row.names(peak_factors),
                     row.names(scent)),]
# Non/metric-multidimensional scaling
scent_nmds <- vegan :: metaMDS(comm = scent, distance = “bray”)
scent_nmds <- as.data.frame(scent_nmds[[ “points” ]])
scent_nmds <- cbind(scent_nmds,
                     colony = peak_factors [[ “colony” ]])
~~~

The results results of the NMDS analysis are outputted to the data frame `scent_nmds` and can be visualised using the package ggplot2 [36].

~~~
# load package ggplot2
library (ggplot2)
# create the plot (see Fig 6)
ggplot(data = scent_nmds, aes(MDS1,MDS2, color = colony)) +
           geom_point () +
           theme_void () +
           scale_color_manual(values = c(“blue”, ”red”)) +
           theme(panel.background = element_rect(colour = “black”,
             size = 1.25, fill = NA),
             aspect.ratio = 1,
             legend.position = “none”)
~~~

The resulting NMDS plot shown in Fig 6 reveals a clear pattern in which seals from the two colonies cluster apart based on their chemical profiles, as shown also by Stoffel et al. [6]. Although a sufficient number of standards were lacking in this example dataset to calculate the internal error rate (as shown below for the three bumblebee datasets), the strength of the overall pattern suggests that the alignment implemented by `GCalignR` is of high quality.

**Fig 6.**
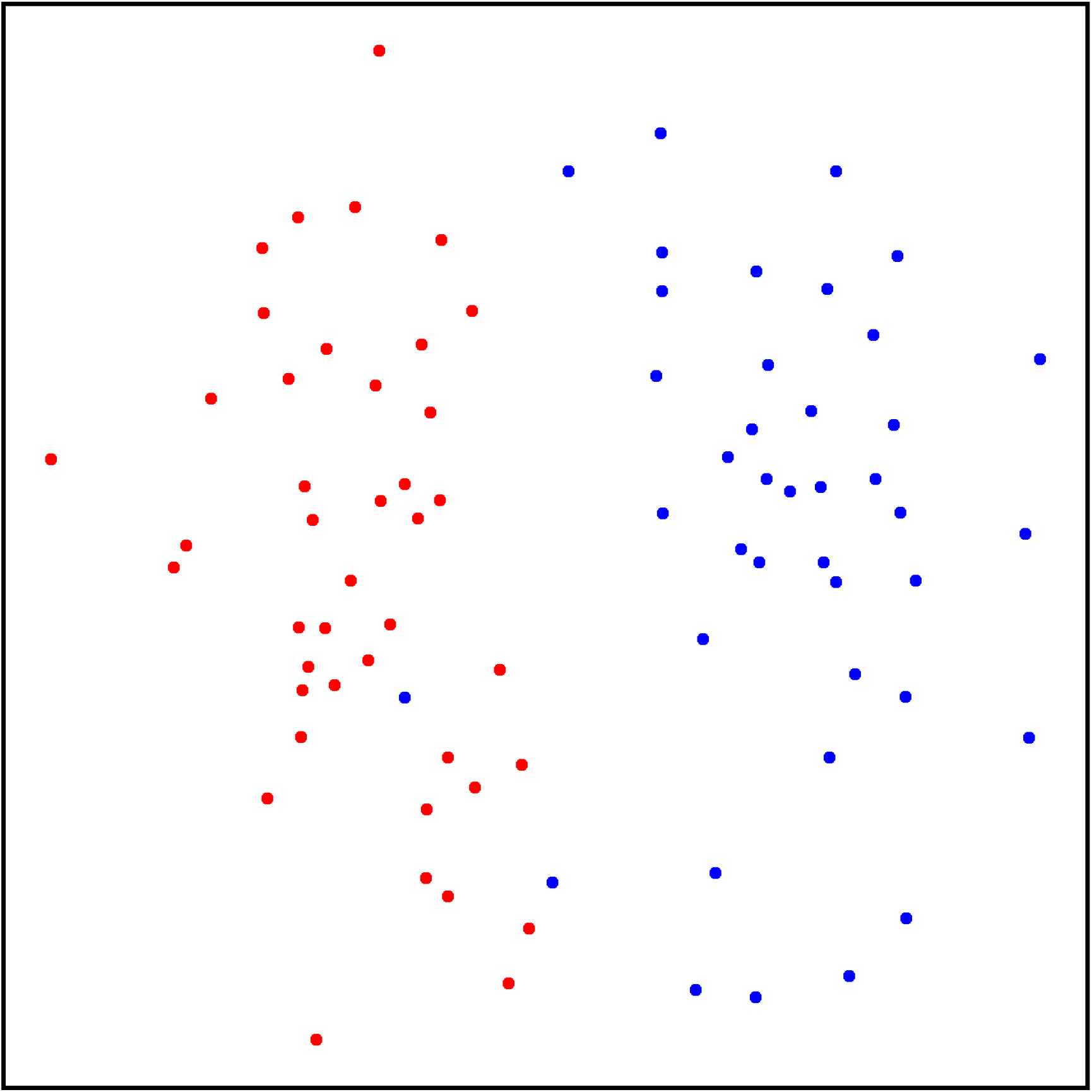
Two-dimensional nonmetric multidimensional scaling plot of chemical data from 41 Antarctic fur seal mother–offspring pairs. Bray–Curtis dissimilarity values were calculated from standardized and log(x+1) transformed abundance data (see main text for details). Individuals from the two different breeding colonies described in Stoffel et al. [6] are shown in blue and red respectively.

### Evaluation the performance of GCalignR

We evaluated the performance of `GCalignR` in comparison to GCALIGNER [23]. For this analysis, we focused on three previously published bumblebee datasets that were published together with the GCALIGNER software [23]. These data are well suited to the evaluation of alignment error rates because subsets of chemicals within each dataset have already been identified using GC-MS [23]. Hence, by focusing on these known substances, we can test how the two alignment programs perform. Furthermore, these datasets allow us to further investigate the performance of `GCalignR` by evaluating how the resulting alignments are influenced by parameter settings.

### Comparison with GCALIGNER

To facilitate comparison of the two programs, we downloaded raw data on cephalic labial gland secretions from three bumblebee species [23] from http://onlinelibrary.wiley.com/wol1/doi/10.1002/jssc.201300388/suppinfo. Each of these datasets included data on both known and unknown substances, the former being defined as those substances that were identified with respect to the NIST database [37]. The three datasets are described in detail by [23]. Briefly, the first dataset comprises 24 *Bombus bimaculatus* individuals characterised for a total of 41 substances, of which 32 are known. The second dataset comprises 20 *B. ephippiatus* individuals characterised for 64 substances, of which 42 are known, and the third dataset comprises 11 *B. flavifrons* individuals characterised for 58 substances, of which 44 are known.

To evaluate the performance of GCALIGNER, we used an existing alignment provided by [23]. For comparison, we then separately aligned each of the full datasets within `GCalignR` as described in detail in S2. We then evaluated each of the resulting alignments by calculating the error rate, based only on known substances, as the ratio of the number of incorrectly assigned retention times to the total number of retention times (Eq (5)).

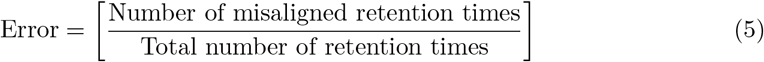

where retention times that were not assigned to the row that defines the mode of a given substance were defined as being misaligned. Fig 7 shows that both programs have low alignment error rates (i.e. below 5%) for all three datasets. The programs performed equally well for one of the species (*B. flavifrons*), but overall `GCalignR` tended to perform slightly better, with lower alignment error rates being obtained for *B. bimaculatus* and *B. ephippiatus*.

**Fig 7.**
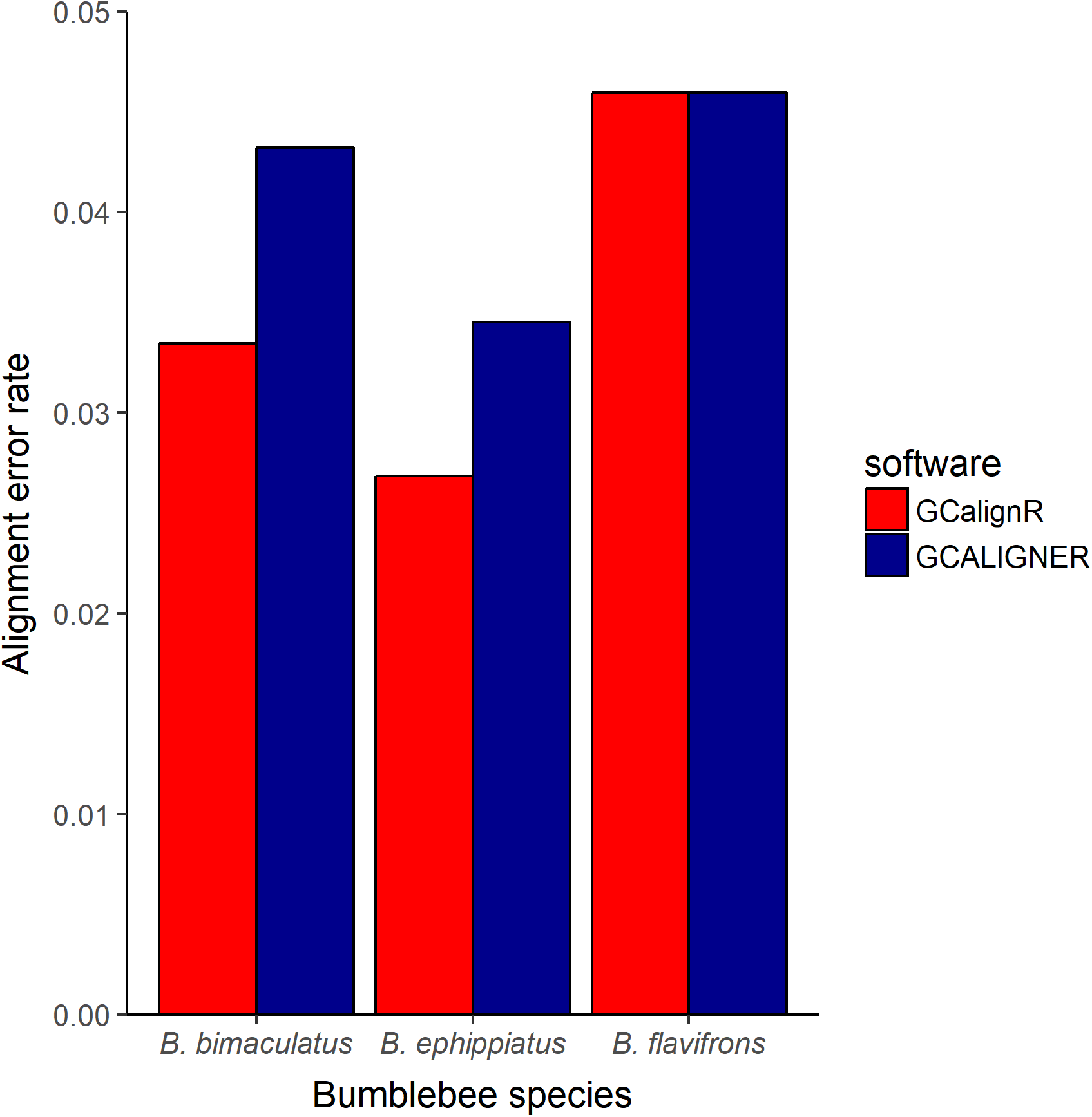
Alignment error rates for three bumblebee datasets using GCalignR and GCALIGNER. Error rates were calculated based only on known substances as described in the main text.

### Effects of parameter values on alignment results

The first step in the alignment procedure accounts for systematic linear shifts in retention times. As most datasets will require relatively modest linear transformations (illustrated by the Antarctic fur seal dataset in Fig 5), the parameter `max_linear_shift` (Table 1), which defines the range that is considered for applying linear shifts (i.e. window size), is unlikely to appreciably affect the alignment results. By contrast, two user-defined parameters need to be chosen with care. Specifically, the parameter `max_diff_peak2mean` determines the variation in retention times that is allowed for sorting peaks into the same row, whereas the parameter textttmin_diff_peak2peak enables rows containing homologous peaks that show larger variation in retention times to be merged (see Material and methods for details and Table 1 for definitions). To investigate the effects of different combinations of these two parameters on alignment error rates, we again used the three bumblebee datasets, calculating the error rate as described above for each conducted alignment. Fig 8 shows that for all three datasets, relatively low alignment error rates were obtained when `max_diff_peak2mean` was low (i.e. around 0.01–0.02 minutes). Error rates gradually increased with larger values of `max_diff_peak2mean`, reflecting the incorrect alignment of non-homologous substances that are relatively similar in their retention times. In general, alignment error rates were relatively insensitive to parameter values of `min_diff_peak2peak` (see Fig 8). Higher error rates were only obtained when `max_diff_peak2mean` was larger than or the same as textttmin_diff_peak2peak, in which case merging of homologous rows is not possible.

**Fig 8.**
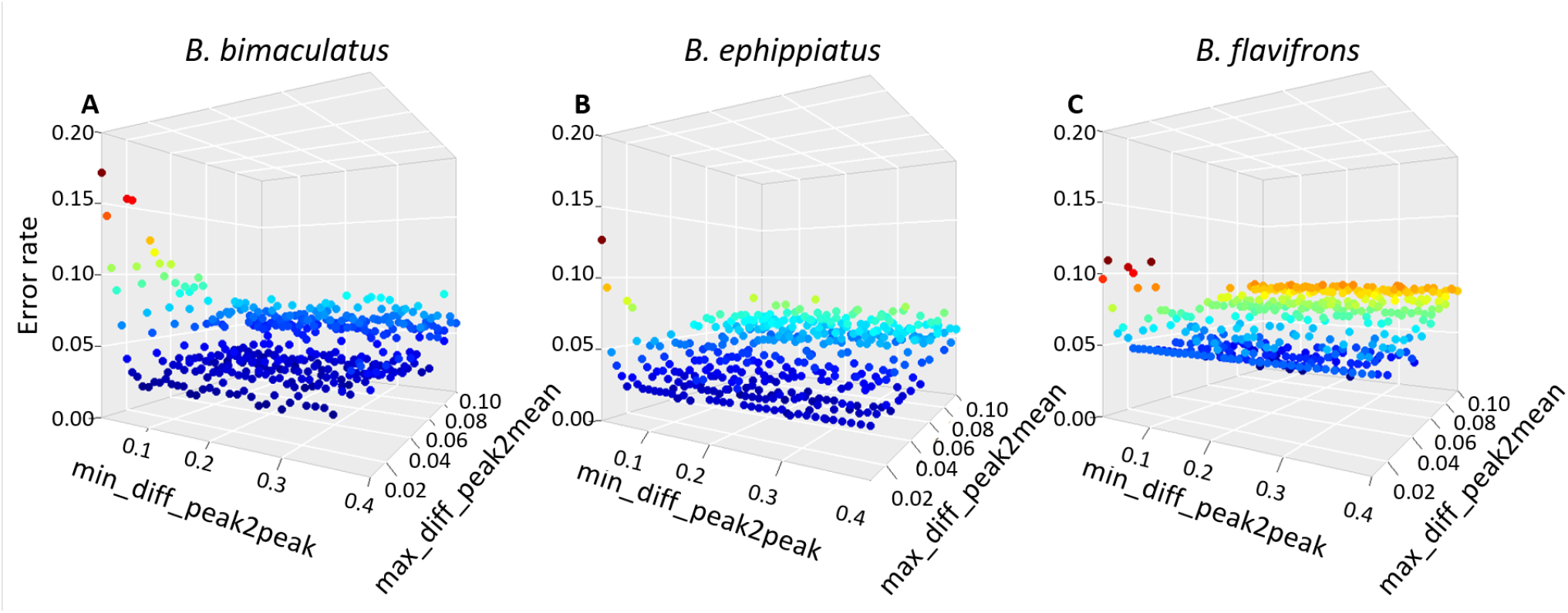
Effects of different parameter combinations on alignment error rates for three bumblebee datasets (see main text for details). Each point shows the alignment error rate for a given combination of `max_diff_peak2mean` and `min_diff_peak2peak`.

### Comparison with parametric time warping

In the fields of proteomics and metabolomics, several methods (usually referred to as ‘time warping’ [18]) for aligning peaks have been developed that aim to transform retention times in such a way that the overlap with the reference sample is maximised [38]. The R package `ptw` [18] implements parametric warping and supports a peak list containing retention times and intensity values for each peak of a sample, making it in principle suitable for aligning GC-FID data. However, parametric time warping of a peak list within `ptw` is based on strictly pairwise comparisons of each sample to a reference [38]. Therefore, the sample and reference should ideally resemble one another and share all peaks [19, 31]. By comparison, `GCalignR` only requires a reference for the first step of the alignment procedure and should therefore be better able to cope with among-individual variability. Additionally, although `ptw` transforms individual peak lists relative to the reference, it does not provide a function to match homologous substances across samples.

In order to evaluate how these differences affect alignment performance, we analysed GC-MS data on cuticular hydrocarbon compounds of 330 European earwigs (*Forficula auricularia*) [39] using both `GCalignR` and `ptw`. This dataset was chosen for two main reasons. First, alignment success can be quantified based on twenty substances of known identity. Second, all of the substances are present in every individual, the only differences being their intensities. Hence, among-individual variability is negligible, which should minimise issues that may arise from samples differing from the reference. As a proxy for alignment success, we compared average deviations in the retention times of homologous peaks in the raw and aligned datasets, with the expectation that effective alignment should reduce retention time deviation.

For this analysis, we downloaded the earwig dataset from https://datadryad.org/resource/doi:10.5061/dryad.73180 [22] and constructed input files for both `GCalignR` and `ptw`. We then aligned this dataset using both packages as detailed in supporting information S2. Following fine-tuning of alignment parameters within `GCalignR`, we obtained twenty substances in the aligned dataset and all of the homologous peaks were matched correctly (i.e. every substance had a retention time deviation of zero). Consequently, `GCalignR` consistently reduced retention time deviation across all substances relative to the raw data (Fig 9). By comparison, parametric time warping resulted in higher deviation in retention times for all but two of the substances (Fig 9). These differences in the performance of the two programs probably reflect differential sensitivity to variation in peak intensities.

**Fig 9.**
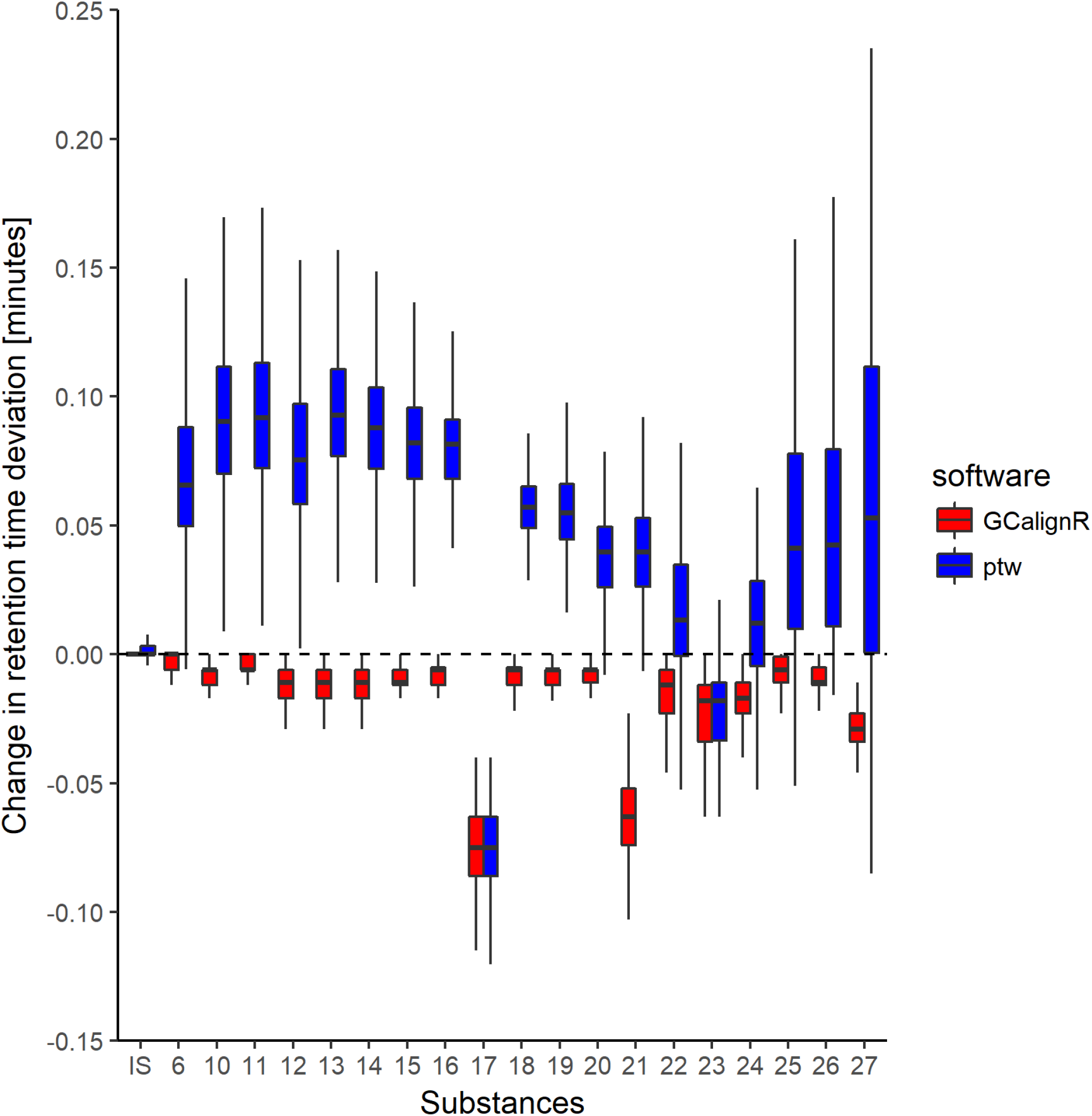
Boxplot showing changes in retention time deviation of twenty homologous substances relative to the raw data after having aligned a dataset of 330 European earwigs within GCalignR and ptw respectively (see main text for details).

## Conclusions

`GCalignR` is primarily intended as a pre-processing tool in the analysis of complex chemical signatures of organisms where overall patterns of chemical similarity are of interest as opposed to specific (i.e. known) chemicals. We have therefore prioritised an objective and fast alignment procedure that is not claimed to be free of error. Nevertheless, our alignment error rate calculations suggest that `GCalignR` performs well with a variety of example datasets. `GCalignR` also implements a suite of diagnostic plots that allow the user to visualise the influence of parameter settings on the resulting alignments, allowing fine-tuning of both the pre-processing and alignment steps (Fig 1). For tutorials and worked examples illustrating the functionalities of `GCalignR`, we refer to the vignettes that are distributed with the package and supporting information S4 and S5.

## Availability

The `GCalignR` release 1.0.0 is available on CRAN and can be installed with recent R versions using the following command:

~~~
install.packages(“GCalignR”)
~~~

We aim to extend the functionalities of GCalignR in future and the most recent release and developmental versions can be accessed on the GitHub repository of the package. Within R, the developmental version can be installed using devtools [40].

~~~
library (devtools)
install_github (“mottensmann/GCalignR”, build_vignettes = TRUE)
~~~

## Supporting information

**S1. A text file.** Details on the bibliographic survey search

**S2. Rmarkdown document.** The R code and accompanying documentation for all analyses presented in this manuscript are provided in an Rmarkdown document file.

**S3. A package vignette.** The vignette ’`GCalignR`: Step by Step’ gives an more detailed introduction into the usage of the package functionalities to tune parameters for aligning peak data.

**S4. A package vignette.** The vignette ’`GCalignR`: How does the Algorithms works?’ gives an introduction into the concepts of the algorithm and illustrates how each step of the alignment procedure alters the outcome based on simple datasets consisting of simulated chromatograms.

**S5. Datasets used to generate the results presented in this manuscript.** This is a compressed zip archive that includes all the raw data that were used to produce the results shown in the manuscript and in S2.

## Acknowledgments

We are grateful to Barbara Caspers and Sarah Golüke for helpful discussions and providing data for testing purposes during the development of the package. This work was funded by a Deutsche Forschungsgemeinschaft standard Grant HO 5122/3-1 (to J.I.H.).

